# Visual and auditory deep learning models capture neural representations of naturalistic social interaction in the superior temporal sulcus

**DOI:** 10.64898/2026.08.13.744446

**Authors:** Itai Peleg, Shiri Almog, Maya Kadushin, Idan Grosbard, Nitzan Guy, Ido Tavor, Galit Yovel

**Author notes:** **Corresponding authors**: Itai Peleg, Sagol School of Neuroscience, Tel Aviv University, Galit Yovel, School of Psychological Sciences & Sagol School of Neuroscience, Tel Aviv University.

## Abstract

The superior temporal sulcus (STS) is selectively responsive to multimodal social interactions. Yet studies so far have relied on pre-defined, simplified stimuli or features to uncover the type of information that drives STS activity. We hypothesized that high-dimensional representations from visual and auditory deep learning models would better predict STS responses to naturalistic social interactions. We used self-supervised visual and auditory deep learning models to extract representations of movie frames and audio, respectively, of a TV series participants watched during fMRI scanning. Voxel-wise encoding models of a joint visual-auditory representation outperformed human-made social-affective annotations in predicting STS. Variance partition further revealed visual-auditory posterior-to-anterior gradient within the STS. To interpret what these models encode, we applied Principal Component Analysis to the encoding model weights. In both the visual and auditory models the first dimension tracked social interaction and peaked in the STS, indicating that social interaction is a dominant dimension of STS representation across both modalities. We conclude that the STS represents naturalistic social interaction in a multimodal manner, integrating visual and auditory information, and that visual and auditory deep learning models capture key representational properties of these responses.

## Introduction

From early infancy, humans develop the ability to readily recognize and understand complex social behavior^1,2^. This ability extends to other species^3,4^, underscoring its evolutionary importance. Understanding social information in humans relies on extracting and integrating dynamic visual and auditory information conveyed by faces, bodies, voices and speech. The superior temporal sulcus (STS), proposed to lie within a third visual pathway specialized for dynamic social perception^5–7^, plays a central role in this process^8^. Previous studies have demonstrated that the STS responds to a range of social and perceptual stimuli across visual and auditory modalities, including biological motion^9^, faces, voices^10–12^, and language^13,14^ and to various visual cues for social interactions such as facing, distance or contingent motion^15–17^.

The variety and complexity of the features that entail social interaction and its multi-modal nature have made it challenging to uncover what type of information is encoded by the STS. Different approaches have been used to address this question. One line of research has measured STS responses to isolated visual and auditory stimuli such as dynamic faces, voices or point light displays of interacting and non-interacting people^14,18,19^. While informative, this approach relies on controlled stimuli, selected by the experimenters, which may not adequately represent the richness of the information that the STS processes during natural social interaction^20,21^. To improve ecological validity, Masson and Isik^22^ measured STS responses during naturalistic movie viewing and used a set of theoretically driven features to characterize each scene including low-level visual features (e.g., motion energy, early-layer DNN activations), as well as social-affective features (e.g., the presence of social interactions, speaking, theory of mind, valence, and arousal). Using these features in voxel-wise encoding models, they found that the STS shows unique selectivity for social interactions. However, these features are defined by the experimenter and are largely simplified. For instance, social interaction is coded as a binary present-or-absent variable, potentially overlooking the complexity of the information that naturalistic stimuli convey.

Deep neural networks (DNNs) offer a data-driven alternative, generating rich, high-dimensional representations of visual and auditory input that are not limited to what experimenters can explicitly define. In the visual domain, deep learning models trained on visual images were shown to predict the response of the visual cortex to images^23–25^. However, these same models do not predict STS activity well. Masson and Isik^22^ found that while a DNN trained on object recognition was the preferred feature across most of the visual brain, the STS was a notable exception, where manually annotated social features dominated instead. This suggests that standard visual DNNs trained to classify static images of object categories fail to capture the information that drives responses in the STS. More recently, using a range of image, video and language deep learning models to predict the fMRI response to short videos clips that include human interactions showed lower predictions in the STS than in visual areas in the ventral stream^26^. Moreover, despite the STS’s preference for dynamic stimuli and its selectivity for language and speech, video-based and large language models showed no advantage in predicting STS responses^26^. Notably, none of these models incorporated auditory input, despite the well-established selectivity of the STS for voices and speech.

Work on the STS and social interaction perception has been mainly focused on the role of visual information (for review see Isik^17^). Much of the STS, particularly the anterior portion, but also parts of the posterior STS, responds strongly to auditory stimuli, especially human voices and speech^27,14,5^. Landsiedel and Koldewyn^19^ did examine social auditory information in the STS but presented auditory stimuli in isolation and primarily focused on a visually localized posterior STS region. Visual and auditory responses in the STS have largely been studied in isolation, using different stimuli and different feature sets. Yet social interaction is inherently multimodal: we simultaneously observe facial expressions, body movements, vocal tone, and speech content. It therefore remains unclear whether the STS encodes a common dimension of social interaction information across modalities or reflects sensitivity to modality-specific features such as faces and voices.

In the current study, we addressed this gap using large-scale self-supervised models trained on large datasets. For the visual domain, we used Contrastive Language-Image Pre-training CLIP^28^, and for the auditory domain, we used Contrastive Language-Audio Pre-training (CLAP^29^), which applies the same contrastive framework to learn joint representations of audio and natural language. Unlike task-specific architectures, these models generate rich multi-dimensional representations of complex visual and auditory input without being optimized for any downstream task. We extracted visual and auditory representations from movie frames and audio of the BBC series Sherlock^30^ and built voxel-wise encoding models to predict fMRI responses of participants (n = 17) who viewed the same stimulus (see Fig. 1 for analysis pipeline). A key challenge of using high-dimensional encoding models is to uncover what dimensions drive their predictions. To interpret these high-dimensional models, we performed principal component analysis (PCA) on the learned encoding weights^31–33^ and visualized their representational geometry, to reveal whether they capture social-semantic structure beyond what hand-crafted annotations encode.

**Fig 1.**
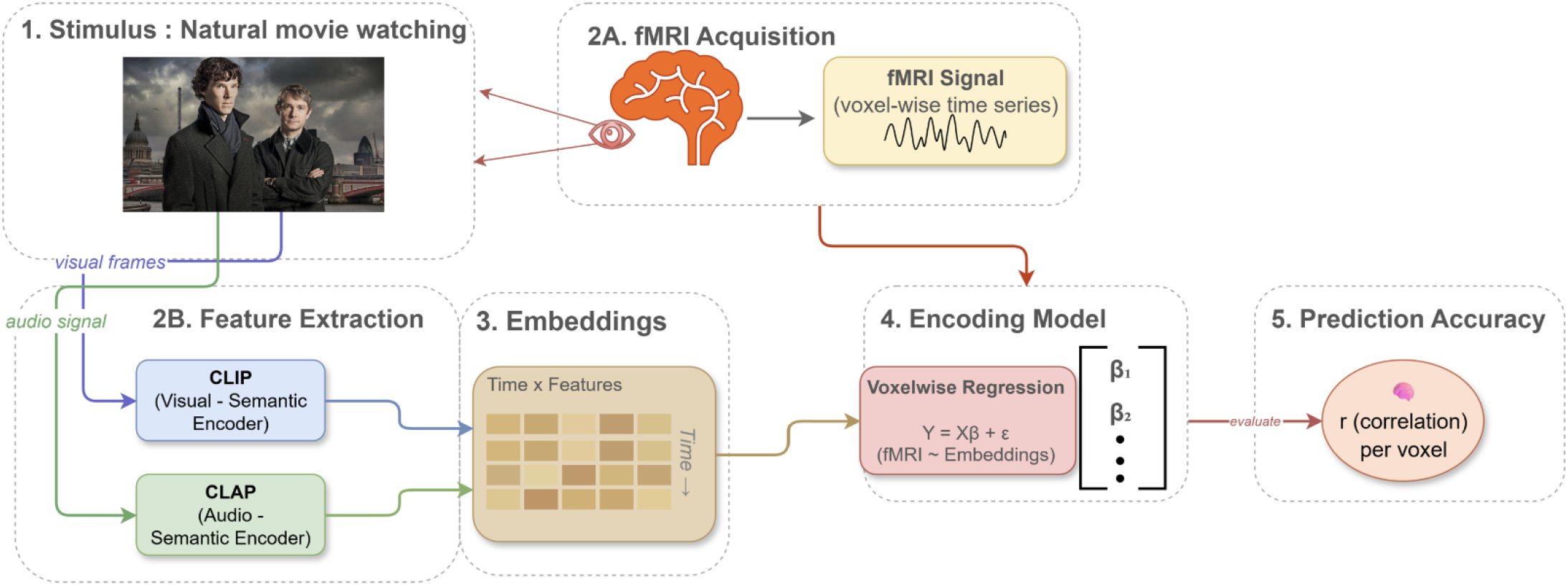
Analysis pipeline from stimulus to voxel-wise prediction via joint CLIP and CLAP embeddings. Participants (N = 17) watched the first episode of the BBC’s Sherlock television series. The stimulus was processed through two channels: (Step 2A) whole-brain fMRI BOLD signals were recorded as voxel-wise time series, and (Step 2B) visual frames and the audio signal were passed through CLIP (ViT-L/14 image encoder)^21^ and CLAP (Contrastive Language–Audio Pretraining)^22^, respectively, to extract feature representations (Step 3). The resulting embeddings were organized into a time points × features matrix, with each row corresponding to one TR (1.5 s) (Step 4). For each voxel, a ridge regression encoding model was fit to predict BOLD responses as a linear function of the DNN embeddings (Step 5). Model performance was evaluated on held-out data by computing the Pearson correlation (r) between predicted and observed BOLD responses for each voxel.

We tested whether a joint CLIP-CLAP model outperforms the manually annotated social-affective model in predicting STS responses and whether the unique contributions of the visual and auditory models dissociate along the posterior-to-anterior axis of the STS. We then asked whether the encoding model weights capture interpretable dimensions of stimulus information. To our knowledge, this is the first study that examines whether auditory deep learning models contribute to STS responses jointly with, and beyond, the predictions of visual deep learning models and whether they extract shared social information. Our results show that the joint CLIP-CLAP model outperforms manually annotated social-affective features in predicting STS responses, with CLIP and CLAP mapping onto distinct posterior and anterior portions of the STS. Analysis of the encoding model weights revealed that social interaction is a dominant dimension that peaks in the STS in both visual and auditory models, indicating that they extract shared social information. Finally, visualization of the representational geometry demonstrated separation of social from non-social content, as well as clustering of semantically similar scenes, revealing rich social-semantic structure in both visual and auditory representations.

## Results

### Joint Vision and Audio DNNs predict neural responses in the STS better than human annotated social-affective features

Our first question was whether multimodal vision-language and auditory-language deep neural networks can serve as effective models for predicting neural responses in the superior temporal sulcus (STS) during naturistic movie viewing. As a first step, we built a joint vision-language and audio-language encoding model. For the visual component, we used the CLIP image encoder (ViT-L/14), which learns a joint latent space for natural language and images^28^. This architecture was chosen because it was shown to produce representations that align well with high-level visual cortex ^33^. For the auditory component, we used CLAP^29^, which similarly learns joint representations of audio and language. We extracted representations from both models using frames and audio from the Sherlock television episode^34^ as input and used these to predict fMRI responses from 17 participants who watched the episode during fMRI scanning^30^. We used the average embeddings of all frames and audio segments within a 1.5 second period, which corresponds with the scanning sampling (TR length) (Figure 1).

We first examined how well the joint CLIP-CLAP model predicts neural responses during natural movie watching. Whole brain analysis has shown high prediction rates in visual and auditory cortex, which peaks in the STS (Figure 2A, Supp Table 1). Next, we asked whether our joint CLIP-CLAP model would capture social information beyond theoretically motivated human annotated social-affective features, which showed high prediction rates of the STS in previous studies^22^. The specific features included in the social-affective model are the presence of social interactions (Binary), speaking (Binary), theory of mind (Mentalization, Category), valence (ordinal), a2nd arousal (ordinal, for more information and the definition of the features see methods and Masson and Isik^22^.

**Fig 2.**
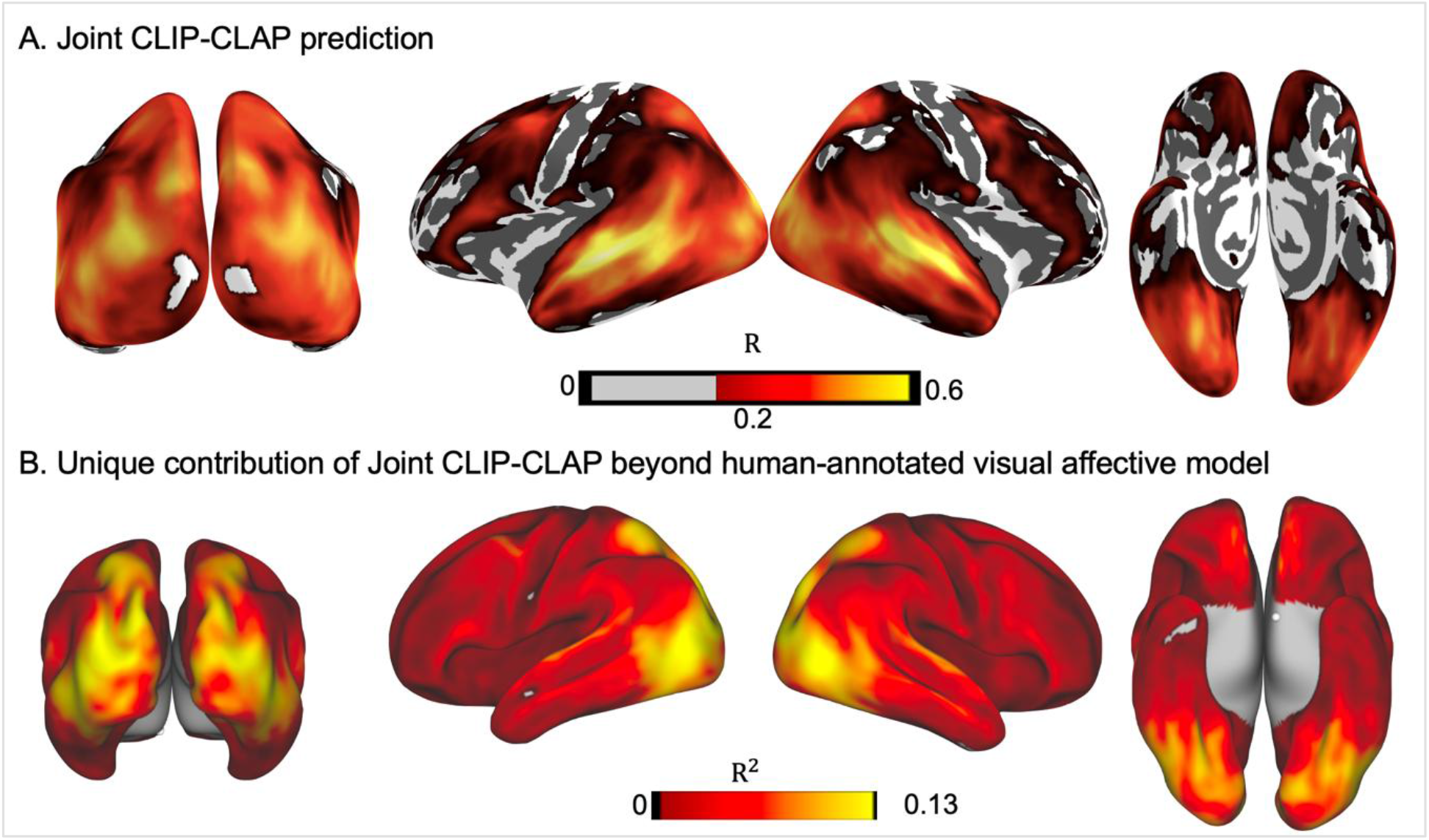
Whole brain neural response predictions by the joint visual-auditory (CLIP-CLAP) model. (A) The CLIP - CLAP model whole brain prediction of neural response during natural movie watching averaged across 17 participants. Pearson correlation smaller than 0.2 (R <0.2) uncolored for visualization (B) Unique variance accounted for by joint CLIP-CLAP model compared to human annotated social affective model.

To assess which model better captured neural responses, we performed variance partitioning with a two-stage statistical approach. First, for each voxel, we compared the unique variance for the joint CLIP-CLAP model over the social-affective model as the difference in R² between their joint contribution relative to the social-affective model contribution [(Unique_CLIP&CLAP_ = R^2^_CLIP&CLAP+SA_ − R^2^_SA_)]. We then tested each voxel whether each model’s unique variance was significantly greater than zero (one-sample t-test, FDR-corrected, p < 0.05). Second, among voxels where at least one model showed significant unique variance, we tested which model explained more unique variance using paired t-tests (FDR-corrected, p < 0.05). This analysis revealed that the joint CLIP-CLAP model accounted significantly more unique variance than the social-affective model (mean ΔR² = 5.8%; range of significant voxels: 0.4%–16.7%), 0.4% marks the smallest significant difference (Fig. 2B).

Next, we performed the same analysis within pre-defined ROIs. These included the posterior and anterior STS, defined using the functional parcellation of Deen^14^, as well as the extrastriate body area (EBA), fusiform face area (FFA), and parahippocampal place area (PPA), which were functionally defined using meta-analytic masks from Neurosynth^35^.

To obtain a cross-subject measure of encoding models’ performance, we computed the mean R² across 150 randomly selected voxels from each ROI and each hemisphere (Fig. 3). The joint CLIP-CLAP model explained significantly more variance than the social-affective model in all ROIs (paired t-tests, *** p < 0.001 for the model comparison, FDR-corrected; see Table S1).

**Figure 3:**
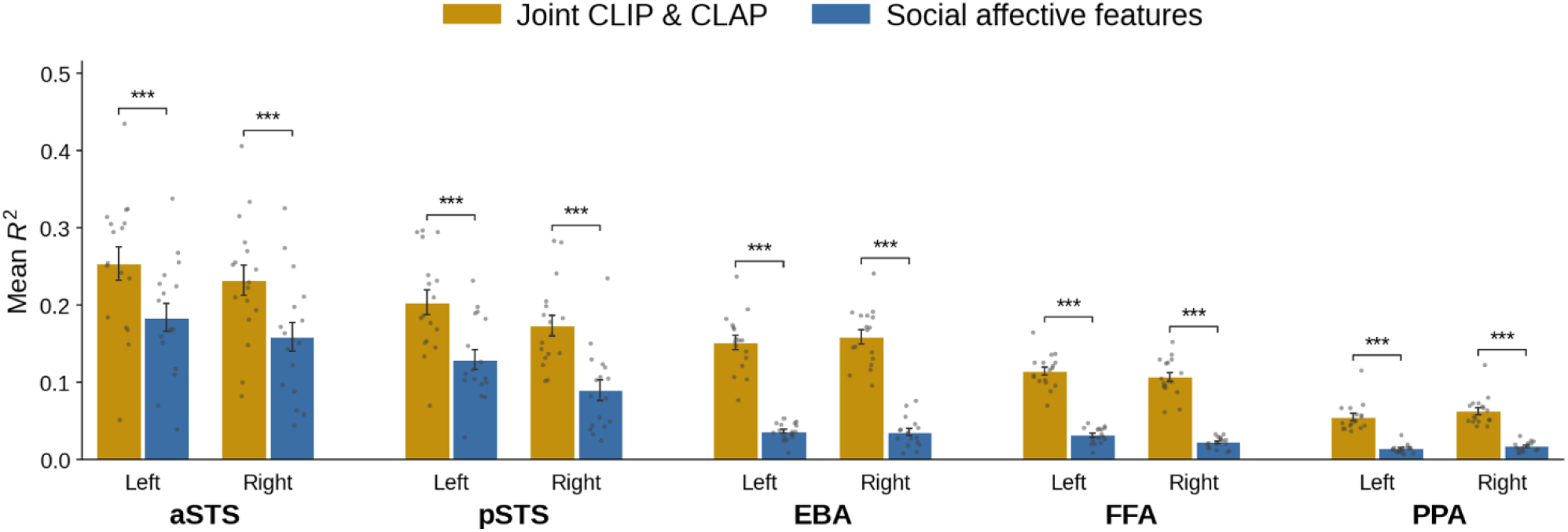
A Joint CLIP-CLAP model better predicts the neural response than social affective features: The mean R^2^ on randomly selected 150 voxels in the STS and category selective areas in the ventral visual cortex is higher for the joint CLIP-CLAP model than human-annotated social-affective features.

### Visual and auditory DNNs capture the processing of social information in the STS

Having established that the joint visual-auditory (CLIP-CLAP) model outperforms the social-affective model in the STS, we next sought to compare each DNN prediction to specific social-affective dimensions. We performed variance partitioning between CLIP and CLAP individually against each of the five features that compose the social-affective model: speaking, social interaction presence, arousal, valence, and mentalization

Preference mapping between CLIP and each individual annotation revealed that CLIP explained substantially more unique variance than any single social-affective feature across nearly all high-level visual cortex (Fig. 4A). For arousal, valence, and mentalization, CLIP’s advantage was prominent, with large unique variance differences (|R²| up to 0.15) spanning visual cortex and extending into temporal regions. The annotation of speaking explained approximately 3% more unique variance than CLIP in the left anterior STS. For social interaction presence, no clear preference emerged in the left STS, while CLIP showed a slight advantage in the right anterior STS.

**Figure 4:**
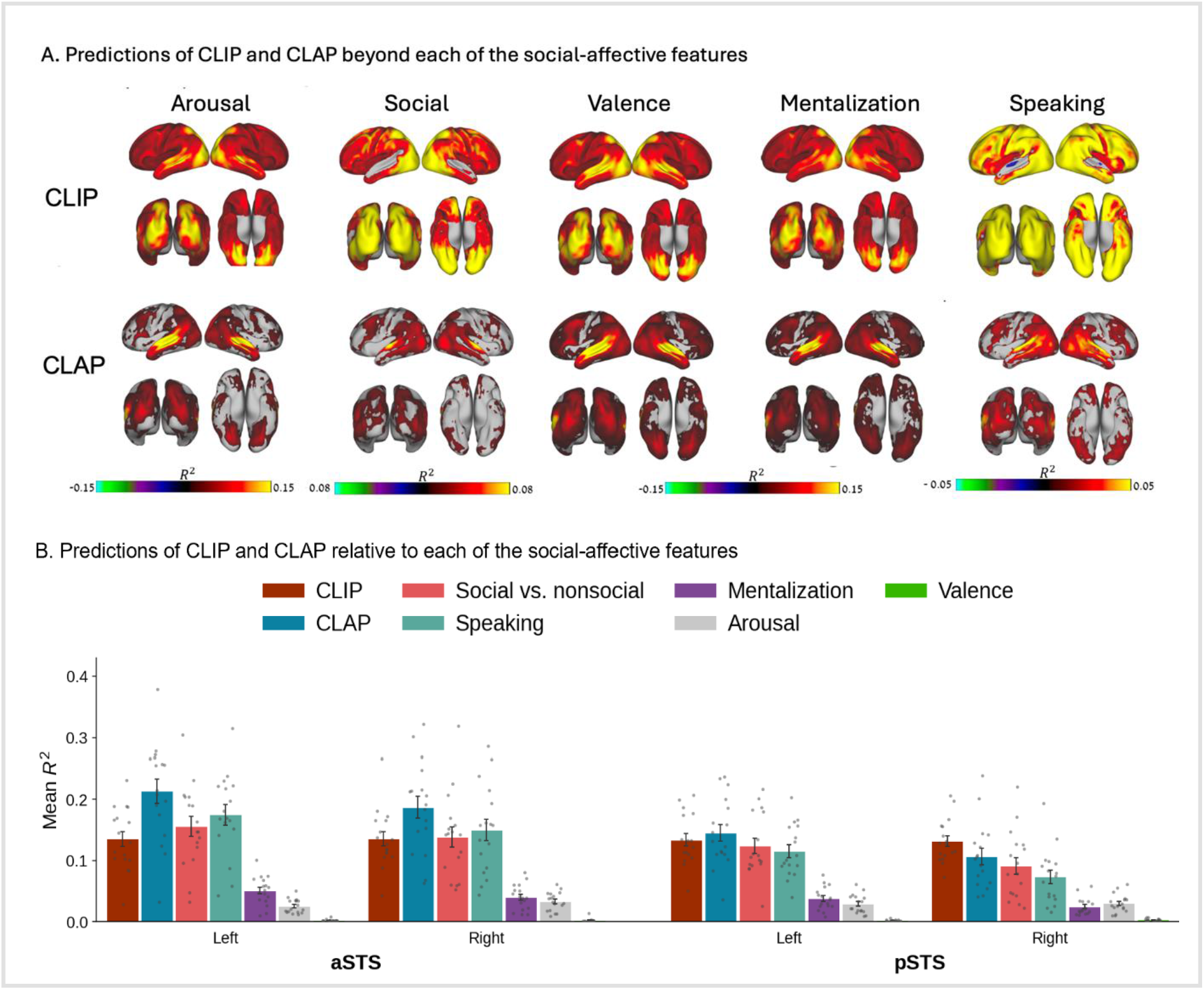
Preference mapping comparing CLIP and CLAP models with each of the social-affective features. (A) Unique variance for CLIP and CLAP versus each of the features of the social-affective model. Positive values (warm colors) indicate voxels where CLIP or CLAP explains more unique variance, negative values indicate social-affective advantage. Only voxels exceeding | |R²| > 0.01 are shown. (B) Model performance measured in mean R² for each ROI and each model and each feature separately, averaged across 17 participants (each dot is a participant).

Preference mapping between CLAP and each individual annotation revealed an advantage for the CLAP across all annotations in temporal cortex (Fig. 4A). For arousal, valence, and mentalization, CLAP’s advantage was broadly distributed, similar to CLIP. For speaking and social interaction presence, CLAP also showed a clear advantage, though with a more selective spatial distribution, notably in anterior and middle STS. Together, these results indicate that CLIP and CLAP capture distinct and complementary information that goes beyond simplified annotations of social information.

### Dissociating visual and auditory contributions to neural prediction

Previous studies have shown that different regions within the STS are sensitive to visual and auditory social information^14^. We therefore examined the unique predictions of the visual (CLIP) and auditory (CLAP) models separately. Whole-brain model comparison between CLIP and CLAP (Fig. 5A) revealed a clear posterior-anterior dissociation. CLIP explained more unique variance throughout visual cortex including the posterior STS, consistent with its role as a visual encoder, while CLAP explained more unique variance in anterior and middle portions of the STS. ROI analysis extended this pattern (Fig. 5B): CLAP predicted significantly more variance than CLIP in the anterior STS bilaterally (left: p < 0.001; right: p < 0.01, FDR-corrected). In the posterior STS, the pattern reversed: CLIP explained significantly more variance than CLAP in the right hemisphere (p < 0.01, FDR-corrected), while no significant difference was observed in the left. Together, these results suggest that CLIP and CLAP capture complementary information, with CLAP encoding auditory and semantic content that extends beyond CLIP’s visual representations, particularly in the anterior STS.

**Figure 5:**
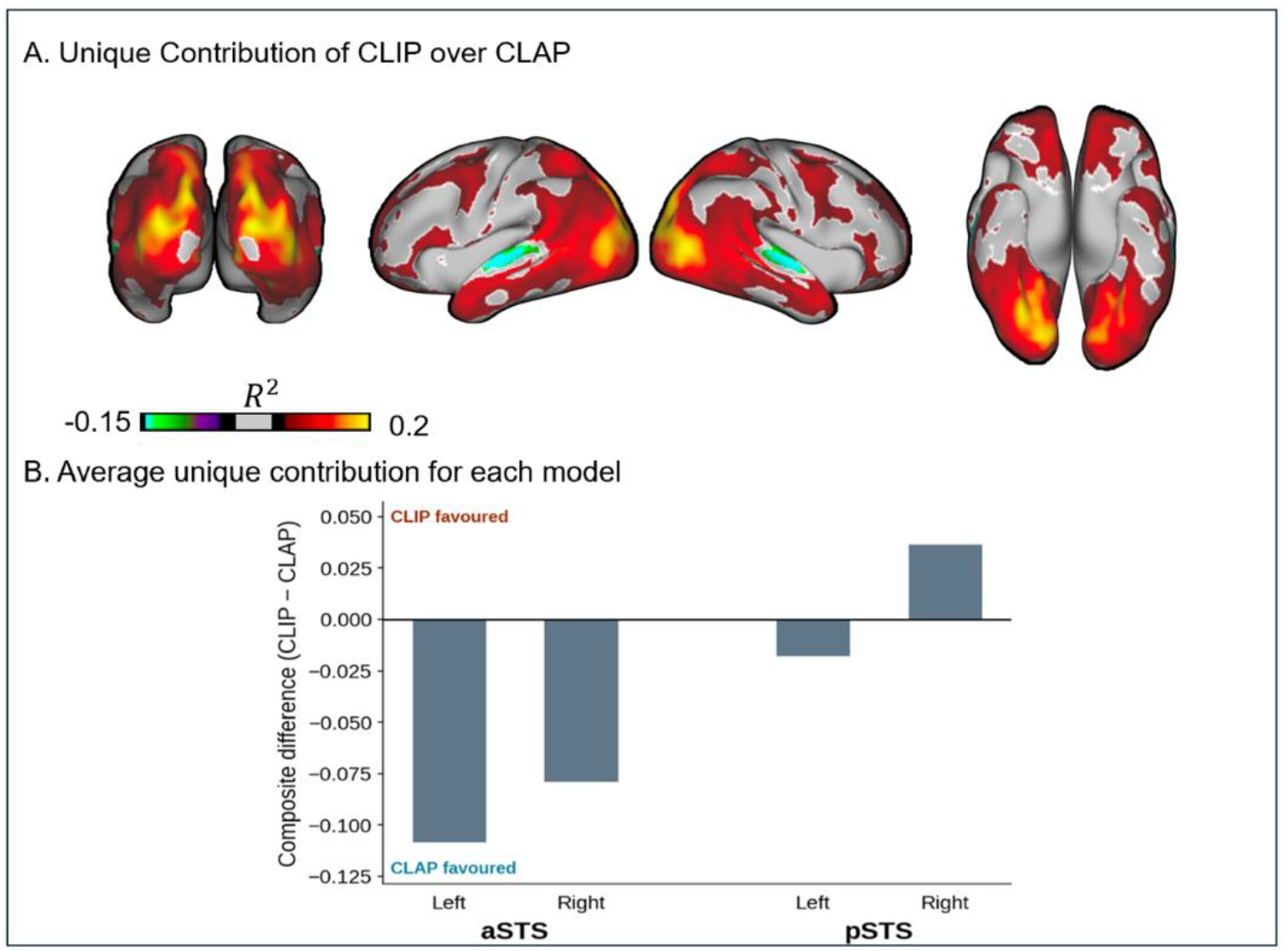
(A) Unique variance for CLIP over CLAP. Positive values (warm colors) are voxels where CLIP explains more unique variance than CLAP, Negative values (cold colors) indicate CLAP advantage. Only voxels exceeding |R²| > 0.05 are shown. (B) Mean unique variance within STS subregions. CLAP (Negative) explains more unique variance in anterior STS bilaterally and left posterior STS, while CLIP (Positive) dominates only in right posterior STS.

### Visual and auditory DNNs represent the processing of visual and auditory social information in the STS

Our analysis shows that joint visual and auditory deep learning models better predict the response of the STS than social-affective features. But what information do they extract from the STS? To explore the semantic dimensions learned by the encoding models, we performed principal component analysis (PCA) on the model weights, following previous work^32,33^. For each model (CLIP and CLAP) we concatenated the learned weight matrices across the predicted voxels per ROI from each of the 17 subjects and voxels from pre-defined ROIs (STS, FFA, PPA), yielding a feature-by-voxel matrix (see Figure S6 for same analysis that includes the EBA). We then ran PCA with voxels as observations, producing principal components (PCs). To obtain a temporal signature for each PC, we projected the per-TR stimulus embeddings onto the PC axes, yielding a PC activation time course for each component. We then correlated these time courses with human annotations from a social-affective feature set^22^ to interpret what each PC represents. Finally, we projected each voxel’s PC score back to its anatomical location and averaged across subjects to visualize the cortical distribution of each PC.Across both the visual (CLIP) and auditory (CLAP) models, the first principal component (PC1) captured information about social interaction. For CLIP, PC1 (explained variance 30.4%) correlated strongly with the presence of social interaction (r = 0.62) and the presence of a face (r = 0.65) (Fig. 6 top right). To test whether PC1 reflects social interaction specifically rather than faces in general, we constructed a face-only vector flagging TRs containing a face but no social interaction. This face-only vector correlated negatively with PC1 (r = -0.23), indicating that the STS does not respond to faces per se but to their appearance in the context of social interaction. Inspection of the PCs scores across ROIs showed that PC1 separates STS from category selective areas FFA, and PPA (Fig. 6, bottom right). Brain projections of CLIP PC1 were localized to lateral temporal cortex including the STS (Fig. 7). Inspection of movie frames at the extreme ends of PC1 is consistent with this interpretation: high-scoring frames depict scenes with multiple interacting characters, while low-scoring frames show environmental shots or individuals with no social interaction present (Fig. 8).

**Fig 6.**
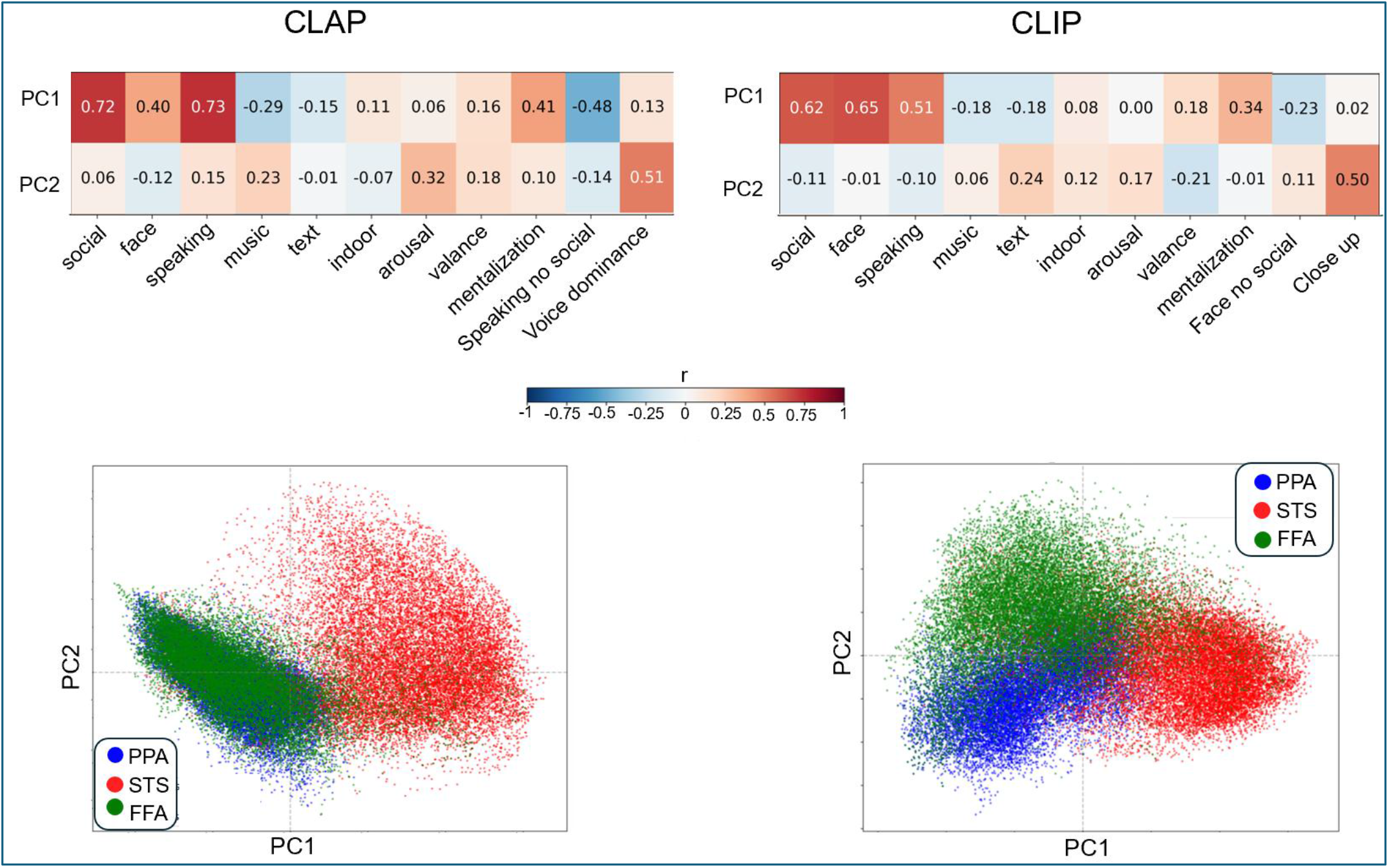
Principal components of the CLIP and CLAP encoding weights capture social-affective-perceptual information in neural responses. (Top) Correlations between the **PC1** activation time course and human-annotated social-affective features, for CLAP (left) and CLIP (right). For CLAP, **PC1** correlated most strongly with social interaction (r = 0.72) and speaking (r = 0.73); for CLIP, **PC1** correlated most strongly with social interaction (r = 0.62) and the presence of a face (r = 0.65). In both models this reflects social interaction specifically rather than faces or voices per se: a face-only vector correlated negatively with CLIP PC1 (r = −0.23), and a speaking-only vector correlated negatively with CLAP PC1 (r = −0.48). **(Bottom)** Each voxel’s loading on PC1 (x-axis) versus PC2 (y-axis), with colors denoting functional ROIs (PPA, STS, FFA). Along PC1, STS voxels separate from the category-selective FFA and PPA in both models, indicating that PC1 distinguishes the STS from ventral category-selective cortex.

**Fig 7.**
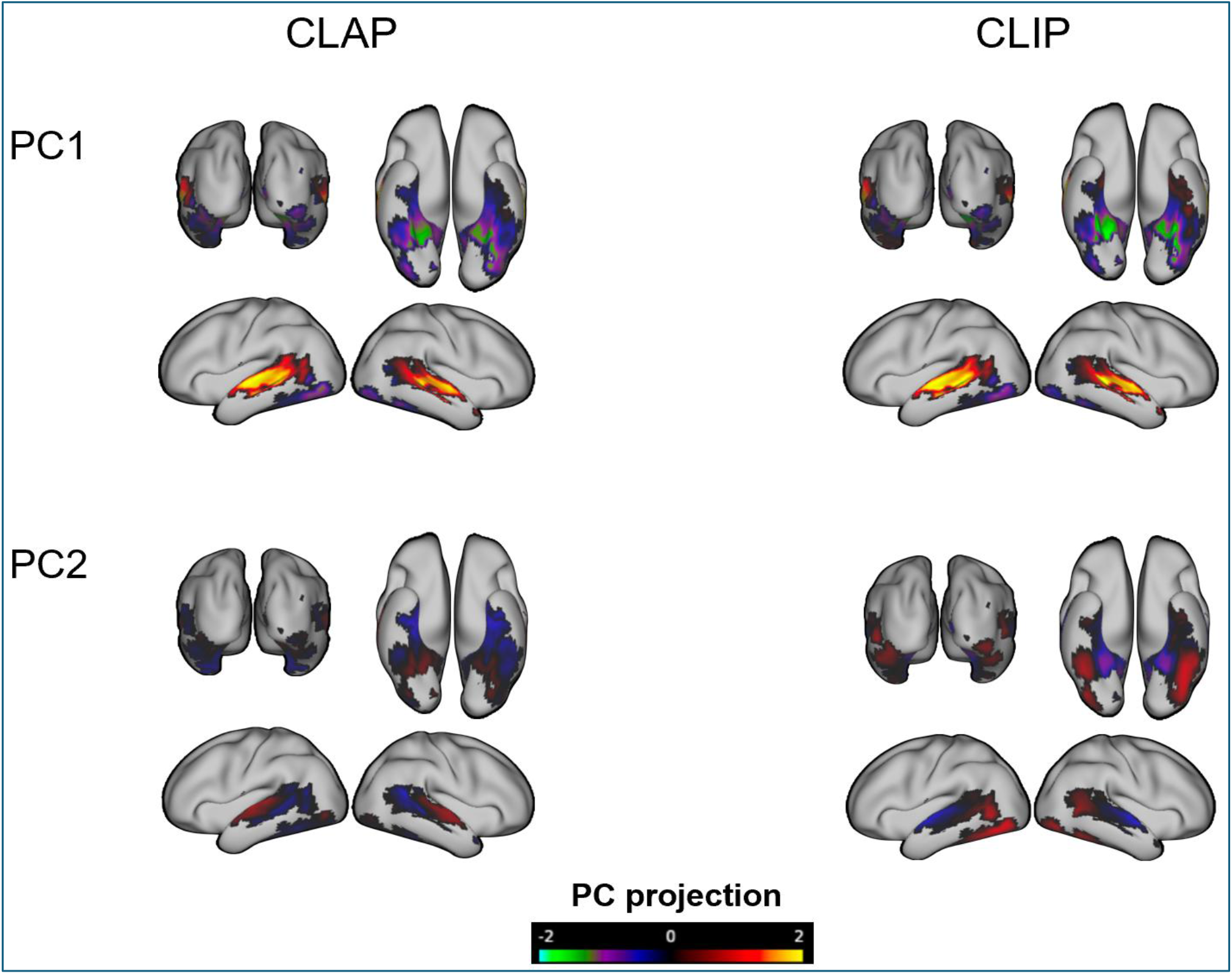
Brain projections of principal components for CLIP and CLAP. PC loading scores projected onto inflated cortical surfaces for CLAP (left) and CLIP (right), for PC1 and PC2. Warm colors indicate positive loadings; cool colors indicate negative loadings (scale: −1.5 to 1.5). **PC1** localizes to bilateral STS in both models with a shared social-interaction dimension. **PC2** dissociates between the two models in both its stimulus correlate and its cortical distribution. In **CLAP**, PC2 correlated with emotional dominance (r = 0.51 with WavLM-derived emotional dominance) and forms an anterior–posterior gradient along the STS. In **CLIP**, PC2 tracks shot scale (r = 0.50 with ShotVL-estimated close-up vs. wide views) and loads onto ventral temporal cortex, including the FFA and posterior STS.

**Fig 8.**
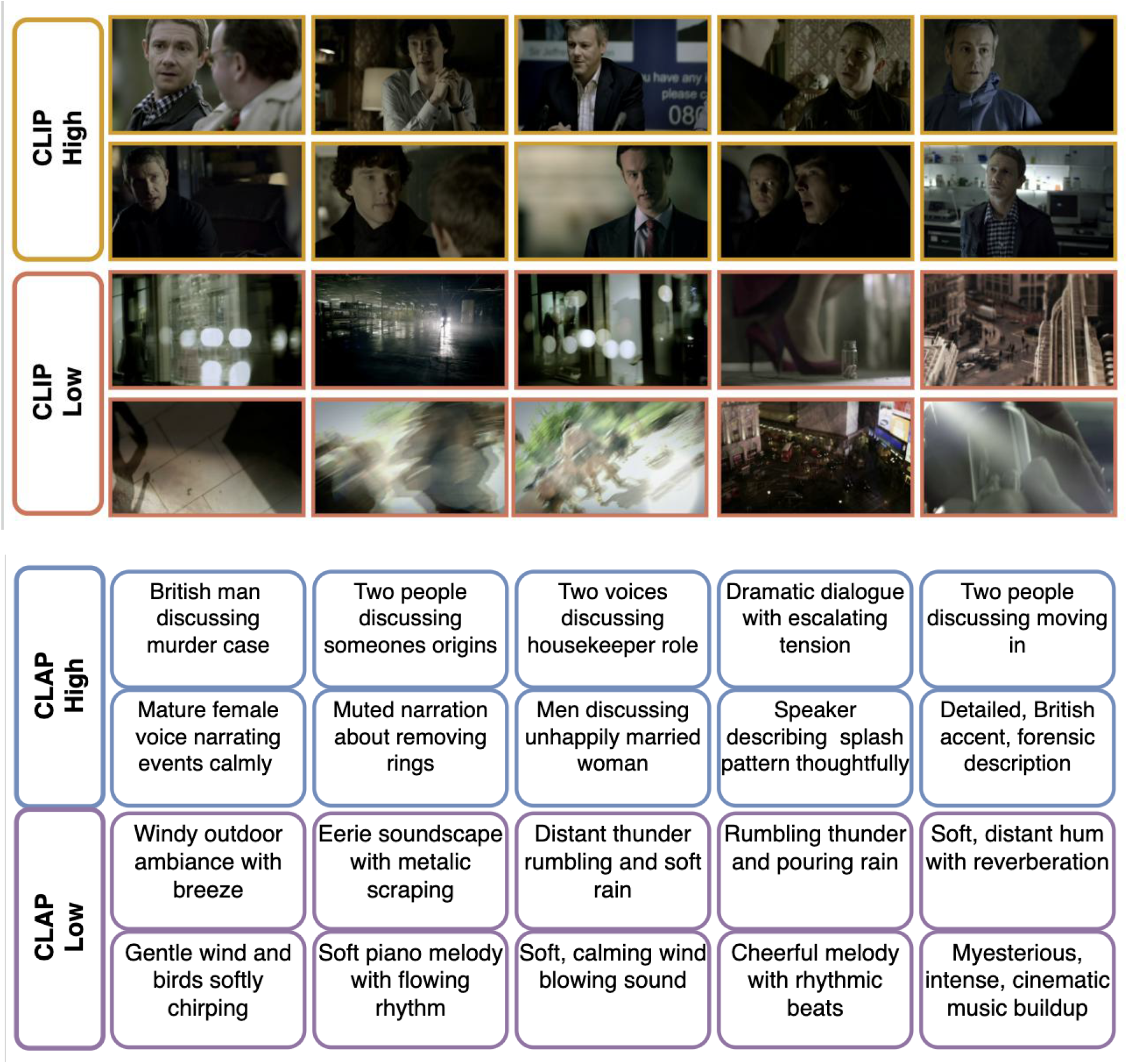
Top and bottom Images and sounds from scenes based on their loading on PC1 in CLIP and CLAP. (Top) CLIP. Highest-scoring and lowest-scoring frames from the Sherlock episode, ranked by their CLIP PC1 activation score. PC scores are assigned to 36 frames; frames shown are randomly selected from each bin. High-PC1 frames consistently depict two or more characters engaged in face-to-face social interaction even when only one of the faces is seen in the image. Low-PC1 frames contain no characters, featuring instead empty environments, outdoor city scenes, and abstract or motion-blurred shots. This contrast directly illustrates the social-interaction dimension captured by CLIP PC1. (Bottom) CLAP. Short descriptions of the highest-scoring and lowest-scoring audio clips, ranked by their CLAP PC1 activation score and summarized with a large audio–language model (GPT-audio). High-PC1 audio-clips consistently contain conversational speech dialogue between two or more speakers and spoken narration (e.g., discussing a murder case, a housekeeper role, or moving in together). Low-PC1 clips contain non-vocal environmental sounds and music with little or no speech, such as wind, rain and thunder, birdsong, and instrumental or cinematic music. This contrast illustrates the social dimension captured by CLAP PC1. See supp Figure for PC2 .

The first principal component (PC1) of CLAP (explained variance 46.4%) showed a similar social-perceptual profile, with strong correlations with social interaction (r = 0.72) and speaking (r = 0.73) (Fig. 6, top left). To assess whether it reflects speaking per se or social related speaking, we computed the correlation of CLAP PC1 with a speaking-only vector (speech without social interaction) and found a negative correlation (r = -0.48), again suggesting that the STS responds to social context rather than the voices alone. Brain projections of CLAP PC1 also localized to lateral superior temporal cortex (Fig. 7).

Inspection of the audio clips at the extreme ends of PC1 is consistent with this interpretation. We described the top-ranked clips using a large audio–language model (GPT-audio^36^), prompting it to summarize each clip in a few words. Clips at the high-scoring extreme consistently contained conversational speech: dialogue between two or more speakers and spoken narration, whereas clips at the low-scoring extreme contained non-vocal environmental sounds and music with little or no speech, such as wind, rain and thunder, birdsong, and instrumental or cinematic music (Fig. 8).

We next examined PC2, which showed dissociation between the ROIs (Fig. S1). For CLIP, inspection of movie frames at the extreme ends of PC2 provides a qualitative characterization: high-PC2 frames tend to feature tight close-up shots, while low-PC2 frames are dominated by wide environmental views (Fig. S1). To quantify this observation, we used ShotVL^37^, a vision-language model trained to classify cinematic shot types, to estimate the shot scale of each frame on a 1 (wide shot) to 5 (close-up) scale. CLIP PC2 (explained variance 6.1%) correlated strongly with shot scale (r = 0.5), indicating that this component captures the distinction between close-up shots of faces and wide environmental views. Brain projections of CLIP PC2 (Fig. 7) loaded on the FFA and nearby ventral temporal cortex (posterior STS), consistent with a shot-scale axis in visual cortex. The top frames driving PC2 (Fig. S1) are tight close-ups, mostly of hands and objects rather than faces, suggesting this axis reflects shot scale (close vs. wide views) rather than face content per se.

For CLAP, listening to audio samples at the extreme ends of PC2 (explained variance 7.6%) revealed a perceptual difference in the emotional quality of the speech. To quantify this, we used a WavLM-based speech-emotion recognition model fine-tuned on the MSP-Podcast corpus^38^ to predict continuous emotion affective attributes from the movie’s audio track, extracting the model’s predicted dominance score per TR. CLAP PC2 correlated with emotional dominance (r = 0.51). Brain projections of CLAP PC2 (Fig. 7) revealed an anterior–posterior gradient along the STS, with positive loadings concentrated anteriorly and negative loadings posteriorly, suggesting a gradient of emotional prosody processing along the temporal lobe that, to the best of our knowledge, has not been reported before.

### Visual and auditory model predictions of the STS do not depend on language supervision

CLIP and CLAP were trained with paired linguistic information, raising the possibility that their predictions of STS responses depend on language-derived semantic supervision. To test this, we ran identical voxel-wise encoding models using two unimodal, self-supervised models trained without any language: DINOv2 (ViT-g/14^39^) for vision and HuBERT^40^ for audio. Each self-supervised model predicted STS responses significantly better than its language-trained counterpart (Fig. S3). DINOv2 outperformed CLIP in the anterior STS (left ΔR² = 0.025; right ΔR² = 0.027) and in the posterior STS (left ΔR² = 0.018; right ΔR² = 0.021), and HuBERT outperformed CLAP in the anterior STS (left ΔR² = 0.026; right ΔR² = 0.022) and in the posterior STS (left ΔR² = 0.017; right ΔR² = 0.017; all paired t-tests p < 0.001, FDR-corrected).

We next asked whether the shared social-interaction dimension identified in CLIP and CLAP will emerge without language supervision as well. Repeating the PCA on the DINOv2 and HuBERT encoding weights recovered the same leading dimension: PC1 correlated with social interaction in both models (DINOv2 r = 0.60; HuBERT r = 0.74), alongside face (DINOv2 r = 0.58) and speaking (HuBERT r = 0.78), and its cortical projection again separated the STS from category-selective face- and scene-responsive regions along PC1. The second component also paralleled the language-trained models, with DINOv2 PC2 tracking shot scale (close-up r = 0.52) and HuBERT PC2 tracking emotional dominance (r = 0.40). The emergence of the same social-interaction dimension in models trained without any language indicates that this dimension reflects structure recoverable from visual and auditory input itself, rather than an artifact of linguistic supervision (Fig. S4).

### Representational geometry of predicted STS responses to social information and scene content

Figure 9 shows the t-SNE visualization of the predicted STS responses for both CLAP and CLIP encoding models, color by social interaction annotation. In both models, the scenes form largely distinct clusters in the two-dimensional space, suggesting that the learned STS representations capture social content from auditory and visual modalities. Notably, semantically coherent scenes tend to group together in this space, for instance scenes from within a vehicle for two distinct clusters. These results provide qualitative evidence that both auditory and visual encoding models learn STS representations that reflect the high-level semantic organization of naturalistic movie content. See Fig. S5 for Visualizations of the predicted STS by the other social-affective annotations.

**Fig 9.**
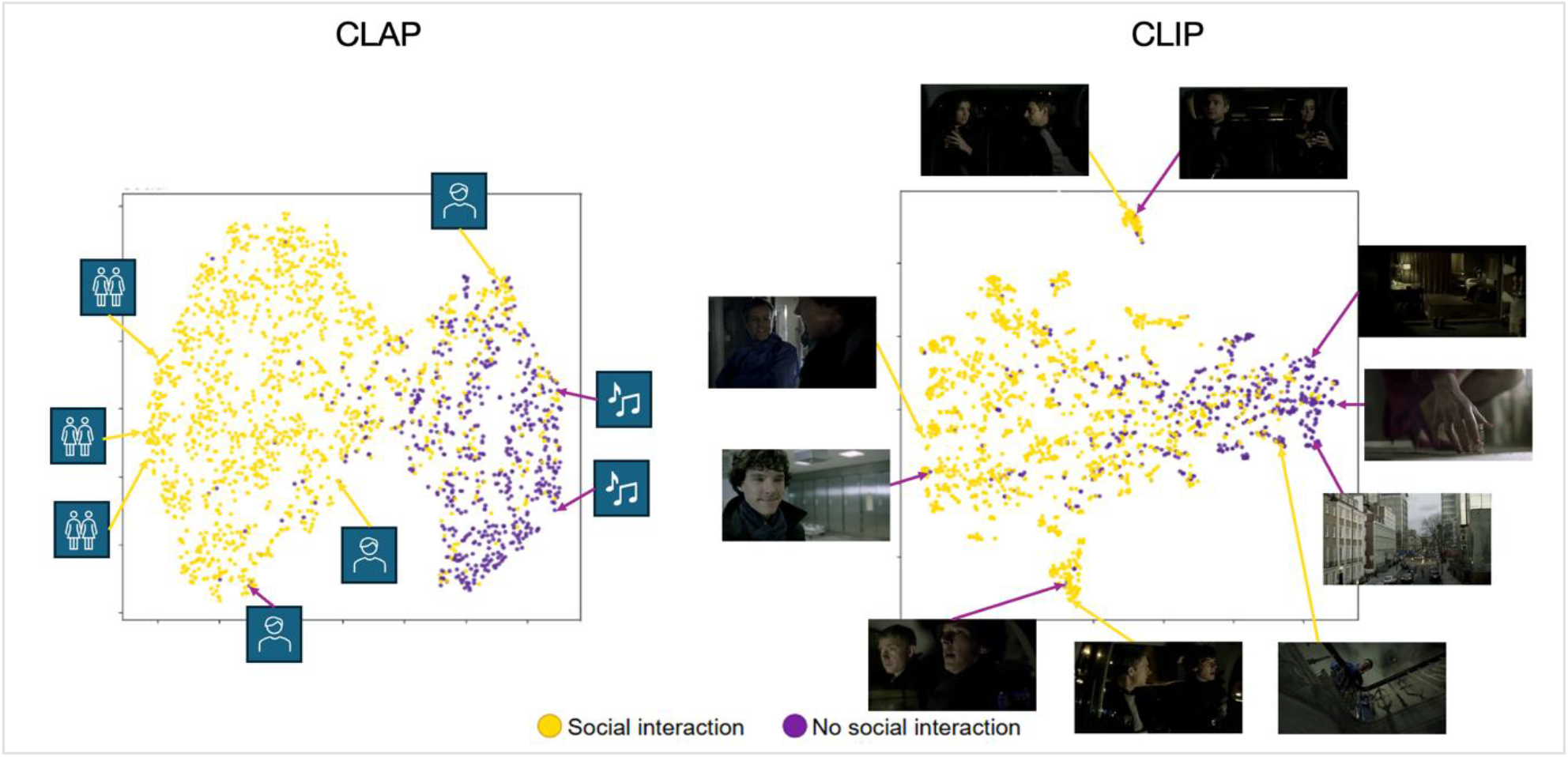
Representational geometry of predicted STS responses. t-SNE visualization of predicted STS activity for CLAP (left) and CLIP (right) encoding models for each movie frame and sound, color-coded by social interaction based on human manual annotations (yellow = social, purple = non-social). The different symbols in CLAP t-SNE are: two people talking, one person and music

## Discussion

What information does the STS represent during natural social perception, and does it integrate visual and auditory content into a shared social dimension? To answer this question, we used the embeddings of joint visual and auditory DNNs, extracted from the frames and audio of natural movies to predict neural responses in participants who watched the TV show Sherlock while undergoing fMRI scanning. This approach revealed three main novel findings. First, joint visual–auditory DNN representations predicted STS responses better than human-annotated social features. Second, auditory representations contributed uniquely and significantly, beyond the visual models that were the focus of previous work^22^. Third, explanatory analysis revealed that the visual and auditory models encoded a shared social-interaction dimension, indicating a representation abstracted from input modality and going beyond prior studies limited to simplified stimuli and experimenter-defined features. This social dimension emerged equally in self-supervised models trained without any language, and therefore does not depend on linguistic supervision.

Previous work on the social representation in the STS has faced a trade-off. Studies using controlled, isolated stimuli such as dynamic faces, voices, or point-light displays gain experimental precision but do not capture the richness of natural social perception^14,15^. Studies using naturalistic stimuli such as movies recover this complexity but have characterized it with human-annotated social features, which are limited to the dimensions an experimenter chooses to define^22,41^. By combining naturalistic movie stimuli with high-dimensional multimodal DNN representations, our approach captures more complex visual and auditory social information than either line of work alone, without limiting that information to experimenter-defined features.

Studies that have used deep learning models to predict responses in high level visual cortex including the STS, have focused on visual models. These studies found that these models better predict the ventral visual cortex than the lateral stream^26^. Our findings show that the auditory model outperforms other features along the STS and more so in the anterior part. This suggests that limiting the investigation of the STS only to its visual features may overlook its inherent multi-modal nature. For example, when looking at a single face in a frame, ignoring auditory information, we may consider this frame as non-social. However, if we hear that the person is listening to another person not seen in the frame that changes the interpretation of the scene to a social scene. This is where the multi-modal information extracted from the STS comes into play to correctly interpret the situation as social. This is consistent with previous reports that show that the STS is responsive to human-related visual-auditory information^14,42^. However, previous studies used isolated face and voice stimuli leaving open the question of whether it extracts shared or distinct social information. Studies that did examine visual-audio interaction in the STS were limited to short video clips of talking faces^43,44^. Our method shows that while the auditory and visual model explained different parts of the variance, they both primarily accounted for complex social information extracted from natural videos, which goes beyond the isolated stimuli used in previous studies.

Interpreting the models revealed a social-interaction dimension shared across the visual and auditory modalities. Although DNNs can model the brain with higher accuracy than was previously possible, their high-dimensional representations make it challenging to uncover what information drives their predictions. To address this, we applied PCA to the encoding model weights and correlated the resulting components with human-annotated social features. The first component, which explained the largest proportion of variance in both the visual (CLIP) and auditory (CLAP) models, correlated strongly with social interaction and projected most heavily onto the STS, indicating that this dimension is represented across both visual and auditory input. The lower components instead captured other dimensions, each projecting onto a functionally matched region: CLAP second principal component corresponded to vocal dominance (r = 0.51) and projected onto anterior STS. CLIP second principal component captured a shot-scale dimension, separating tight close-ups from wide environmental views, and projected onto ventral temporal and lateral occipital cortex.

Visualizing the predicted STS responses with t-SNE offered a complementary, geometry-level view of this organization. Scenes separated primarily by social content, with socially interactive and non-interactive scenes forming largely distinct clusters in both the visual and auditory models. Beyond this social axis, semantically coherent scenes are grouped together, with visually distinct contexts such as conversations inside a car or certain street scenes forming separate clusters. These representations demonstrate that beyond the presence or absence of social interaction, the STS also encodes the particular content of the scene.

To conclude, our findings establish that the STS represents naturalistic social interaction as a high-level dimension that is shared across vision and audition. Using representations from self-supervised visual and auditory deep learning models, we revealed that their leading dimension, in both the visual and auditory models, tracked social interaction and localized to the STS, suggesting a common social representation rather than modality-specific tuning to faces or voices. Some questions remain open. Because our interpretation of the model dimensions relied on a limited, largely binary annotation set, an important next step is to develop richer, graded annotations of social interaction that can capture the continuous structure these models appear to encode. The novel prosodic–emotional gradient revealed by the auditory components leaves an opening for more direct investigation. More broadly, these findings are based on a single naturalistic film and should be tested for generalizability across other stimuli and paradigms. The same approach, combining rich multimodal representations with naturalistic neural data could be extended beyond the STS to characterize how other high-level regions represent complex natural information.

## Method

### Participants and fMRI Data

We analyzed publicly available fMRI data from the Sherlock dataset^30^. In this study, 17 participants (10 male) watched the first episode of the BBC’s Sherlock television series (“A Study in Pink,” 2010^34^; duration ≈ 48 min) in the MRI scanner across two sessions (23 and 25 min each). None of the participants had watched the Sherlock series. The study was approved by the Princeton University Institutional Review Board, and all participants provided written informed consent.

### fMRI Data Acquisition and Preprocessing

Functional images were acquired on a 3T Siemens Skyra scanner using a 20-channel head coil. Whole-brain images (27 slices; voxel size = 4 × 3 × 3 mm³) were collected with a T2*-weighted echo-planar imaging (EPI) sequence (repetition time [TR] = 1,500 ms; echo time [TE] = 28 ms; flip angle = 64°; field of view = 192 × 192 mm).

Preprocessing was performed by the original authors^30^ and included slice-timing correction, motion correction, linear detrending, temporal high-pass filtering (140 s cutoff), spatial normalization to Montreal Neurological Institute (MNI) space with resampling to 3 × 3 × 3 mm³ isotropic voxels, and spatial smoothing with a 6-mm full-width-at-half-maximum (FWHM) Gaussian kernel. Timeseries blood oxygenation level-dependent (BOLD) signals were z-scored across time at every voxel. To account for the hemodynamic response delay, the fMRI data were shifted by 4.5 s (3 TRs) relative to stimulus onset.

### Stimulus Material

The stimulus consisted of the first episode of the Sherlock BBC television series (“A Study in Pink,” 2010)^34^, a 48-minute crime drama involving rich social interactions, dialogue, and dynamic visual scenes. The movie contains extensive social content, including face-to-face conversations, theory of mind inferences, and emotional expressions, making it well-suited for studying social information processing. The episode was divided into 1,924 timepoints (TRs), each corresponding to 1.5 s of movie content.

### Visual Features using CLIP: Contrastive Language–Image Pretraining

For the visual modality, we used the image encoder from CLIP (Contrastive Language–Image Pretraining^28^) with a ViT-L/14 (Vision Transformer, Large, patch size 14) backbone. CLIP learns a joint latent space for images and natural language by training separate image and text encoders on large-scale image–caption pairs using a contrastive objective. The model was selected because prior work has demonstrated that CLIP representations align well with high-level visual cortex^33^. We extracted last-layer image embeddings (dimensionality: 768) from the CLIP visual encoder for each video frame.

Video frames were extracted from the Sherlock episode. For each TR (1.5 s), all frames within that temporal window were passed through the CLIP ViT-L/14 image encoder, and the resulting embeddings were averaged to produce a single visual feature vector per TR.

### Auditory Features using CLAP: Contrastive Language–Audio Pretraining

For the auditory modality, we used the audio encoder from CLAP (Contrastive Language–Audio Pretraining^29^), which learns joint representations of audio and natural language using a contrastive learning framework analogous to CLIP. CLAP maps audio waveforms and text descriptions into a shared embedding space by training an audio encoder^45^ paired with a text encoder ^46^ on the LAION-Audio-630K dataset^29^. We extracted last-layer audio embeddings from the CLAP audio encoder for the movie’s soundtrack.

Audio segments corresponding to each 1.5-s TR were extracted from the movie’s soundtrack and passed through the CLAP audio encoder. This produced a single auditory feature vector per TR.

### Unimodal self-supervised control models (DINOv2 and HuBERT)

To test whether the encoding-model predictions depend on language supervision, we repeated the feature-extraction and encoding pipeline using two unimodal, self-supervised models trained without any paired language. For the visual modality, we used DINOv2 with a ViT-g/14 (giant) backbone^39^, extracting last-layer image embeddings for each video frame and averaging all frames within each 1.5-s TR to produce a single visual feature vector per TR, mirroring the CLIP procedure. For the auditory modality, we used HuBERT^40^, extracting last-layer (768-dimensional) representations from the movie soundtrack and mean-pooling the frame-level features across time within each 1.5-s TR to yield a single auditory feature vector per TR, mirroring the CLAP procedure. These features were used in identical voxelwise encoding models.

### Social-Affective human-generated annotations

We used human-generated annotations from Masson and Isik^22^ of five social-affective features: the presence of social interactions (binary: presence = 1, absence = 0), whether an agent was speaking (binary), whether a character was engaging in theory of mind or mentalization (binary), perceived valence, and arousal. Social interaction (binary), and speaking (binary) labels were generated by two independent raters and showed high inter-rater agreement (r = 0.92 for social interactions in the Sherlock dataset^22^). Valence and arousal ratings were obtained from a separate group of 113 participants who rated 4.5-s video segments on a 1–9 Likert scale^47^. For binary features, annotations were averaged across raters and merged into 1.5-s segments (corresponding to TRs) to align with the fMRI temporal resolution^22^.

### Voxel-wise Encoding Models

We constructed voxel-wise encoding models to predict fMRI responses from CLIP and CLAP representations of the video frames and audio soundtrack, respectively. For each voxel, BOLD responses were modeled as a linear function of the input feature space using ridge regression^48^. Prior to model fitting, all feature vectors were L2-normalized along the feature dimension.

We built three encoding models: (1) a CLIP model using the CLIP image encoder embeddings, (2) a CLAP model using the CLAP audio encoder embeddings, and (3) a joint CLIP–CLAP model using concatenated CLIP and CLAP embeddings. In addition, we built a social-affective encoding model using the five human-annotated social-affective features described above^22^.

Data were split into a training set (80%) and a held-out test set (20%). The ridge regularization parameter was selected from 10 logarithmically spaced values between 10 and 10,000 using 5-fold cross-validation within the training set. The regularization parameter yielding the highest mean R² across folds was used to fit the final model on the full training set. Model performance was evaluated on the held-out test data by computing the Pearson correlation (r) between predicted and observed BOLD responses for each voxel. See Figure 1 for illustration of the analysis pipeline.

### Variance Partitioning

To assess the unique contribution of each model to the prediction of neural responses, we performed variance partitioning analyses^49^. For each voxel and participant, we computed the squared Pearson correlation (r²) between predicted and observed BOLD responses for each encoding model. Unique variance for a model of interest was then computed as: *U(A) = R²(A&B) − R²(B)* where *R²(A&B)* is the variance explained by a joint model with concatenated features from models A and B, and *R²(B)* is the variance explained by model B alone. This approach isolates the variance uniquely explained by model A, above and beyond what model B captures.

We applied this framework to several model comparisons: (a) the joint CLIP–CLAP model versus the social-affective model; (b) CLIP versus CLAP; and (c) each DNN (CLIP or CLAP) versus each individual social-affective annotation (social interaction, speaking, mentalization, valence, and arousal).

Statistical inference was conducted using a two-stage approach. In the first stage, for each voxel, we tested whether each model’s unique variance was significantly greater than zero using a one-sided one-sample t-test across participants. In the second stage, among voxels where at least one model showed significant unique variance, we compared unique variance between the two models using two-sided paired-sample t-tests, with preference assigned to the model with the greater mean unique variance. All p-values were corrected for multiple comparisons using the Benjamini–Hochberg FDR procedure^50^ at α = 0.05.

### Regions of Interest

We defined several regions of interest (ROIs) for targeted analyses. Face-selective, body-selective and place-selective ROIs in the ventral visual cortex were derived from the Neurosynth meta-analytic database^35^ using the terms “face”, “body” and “place”, respectively. A bilateral STS mask was similarly obtained from Neurosynth using the term “superior temporal sulcus.” All Neurosynth association test maps were binarized to create ROI masks (see Supplementary Materials for URLs and map details). For finer-grained analyses of the STS, we additionally divided the STS into posterior (pSTS) and anterior (aSTS) subdivisions bilaterally, based on the functional parcellation of Deen^14^.

### Principal Component Analysis

To uncover the semantic dimensions captured by each encoding model, we performed principal component analysis (PCA) on the learned encoding model weight matrices, following approaches from previous work^32,33^. For each DNN (CLIP and CLAP), we selected the voxels from each ROI for each participant. The corresponding weight vectors were concatenated across all ROIs and all 17 participants along the voxel dimension, yielding a matrix of size F × V, where F is the embedding dimensionality and V is the total number of voxels. This matrix was mean centered along the voxel dimension (subtracting each feature’s mean across voxels), and PCA was applied to the transposed matrix, treating voxels as observations and features as variables.

To interpret the PCs, we correlated the projection of each stimulus timepoint onto each PC with the human-generated social-affective annotation^22^. This revealed which social-semantic dimensions each PC captured. To visualize the cortical distribution of each PC, each voxel’s PCA score was mapped back to its MNI coordinate. In addition, we also explored the top of 200 positive and negative frames of each PC of based on the predictions of the visual and auditory models to examine the content of the visual and auditory information that they convey, respectively.

### Shot-scale estimation (ShotVL)

To characterize the visual dimension captured by CLIP PC2, we estimated the cinematic shot scale of each frame using ShotVL (ShotVL-3B)^37^, a vision–language model fine-tuned for cinematic understanding. For each TR, eight frames were sampled evenly across the 1.5-s window and passed to the model, which was prompted to classify shot scale on a five-point scale (1 = extreme long shot to 5 = extreme close-up). Scores were averaged within each TR to yield a single shot-scale value per TR, which was then correlated with the PC2 time course.

### Speech-emotion estimation (WavLM)

To characterize the auditory dimension captured by CLAP PC2, we estimated continuous emotional attributes from the movie soundtrack using a WavLM-based speech-emotion recognition model fine-tuned on the MSP-Podcast corpus^38^ (the Odyssey 2024 multi-attribute baseline model), which predicts arousal, dominance, and valence^51^. The soundtrack was converted to mono and resampled to 16 kHz, then divided into consecutive 3-s segments. Each segment was normalized using the model’s training statistics and passed through the model to obtain arousal, dominance, and valence predictions. Because each 3-s segment spans two 1.5-s TRs, per-segment predictions were assigned to both corresponding TRs to align with the fMRI sampling rate. The resulting per-TR dominance scores were correlated with the CLAP PC2 time course.

### Audio description with a large audio–language model (GPT-audio)

To qualitatively characterize the auditory content at the extremes of the CLAP principal components, we used OpenAI’s gpt-audio model^36^. Audio clips corresponding to the highest- and lowest-scoring TRs along each PC were submitted to the model via the Chat Completions API. For individual clips, the model was prompted to summarize each clip in a few words. Model outputs were used solely for qualitative interpretation and did not enter any quantitative analysis.

### Visualization of the geometric representation of the visual/auditory content of movie frames

To examine how the predicted neural activity in the STS organizes along social and non-social dimensions, we applied t-distributed Stochastic Neighbor Embedding (t-SNE^52^) to the predicted BOLD responses in the STS. For each encoding model (CLIP and CLAP), we computed predicted STS responses for each TR yielding a matrix of size T × V, where T is the number of TRs and V is the number of STS voxels across all participants. This matrix was projected into a two-dimensional space using t-SNE (perplexity = 30), where each point corresponds to a single TR. TRs were then color-coded according to the social-affective feature^22^. This approach allows us to directly visualize whether the geometry of predicted STS responses naturally separates social from non-social movie content, without imposing any explicit supervision on the dimensionality reduction.

## Supplementary material

**Supp Table 1.** Mean R² ± standard deviation across 17 participants for the joint CLIP-CLAP model and the social-affective model, reported separately for left and right hemispheres across five ROIs (posterior STS, anterior STS, FFA, PPA, EBA). *** indicates p < 0.001, FDR-corrected paired t-test comparing the two models within each ROI. (see Figure2)

|  | Joint CLIP-CLAP |  | Social-Affective model |  |
| --- | --- | --- | --- | --- |
|  | Left | Right | Left | Right |
| Post. STS | $0.2 \pm 0.06$ *** | $0.17 \pm 0.05$ *** | $0.13 \pm 0.05$ | $0.09 \pm 0.05$ |
| Ant. STS | $0.25 \pm 0.08$ *** | $0.23 \pm 0.08$ *** | $0.18 \pm 0.07$ | $0.16 \pm 0.07$ |
| FFA | $0.10 \pm 0.02$ *** | $0.1 \pm 0.02$ *** | $0.03 \pm 0.01$ | $0.02 \pm 0.01$ |
| PPA | $0.04 \pm 0.01$ *** | $0.05 \pm 0.01$ *** | $0.015 \pm 0.005$ | $0.02 \pm 0.005$ |
| EBA | $0.15 \pm 0.04$ *** | $0.16 \pm 0.04$ *** | $0.03 \pm 0.01$ | $0.03 \pm 0.01$ |

**Fig S1:**
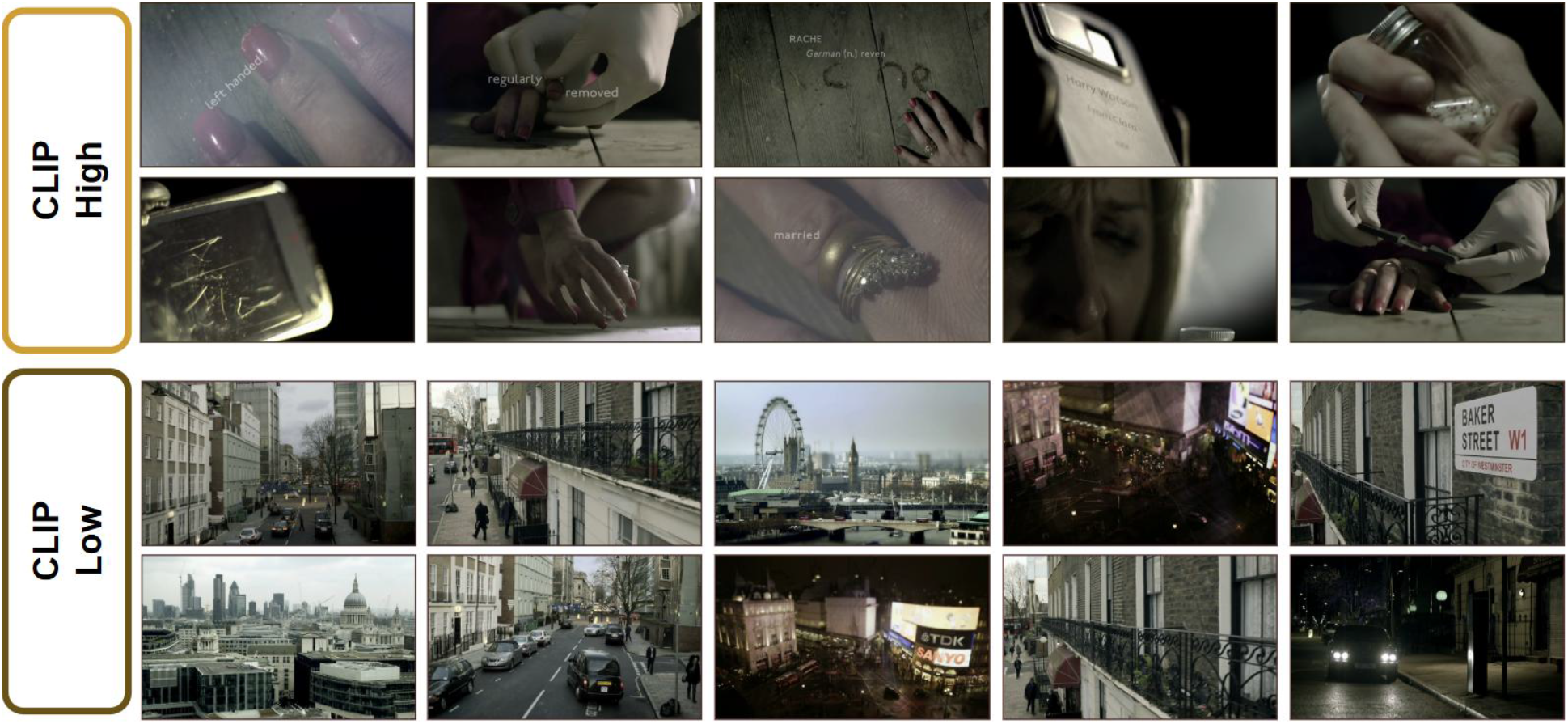
Images from scenes that scored maximum and minimum value for PC2 of CLIP predictions.

**Fig S2:**
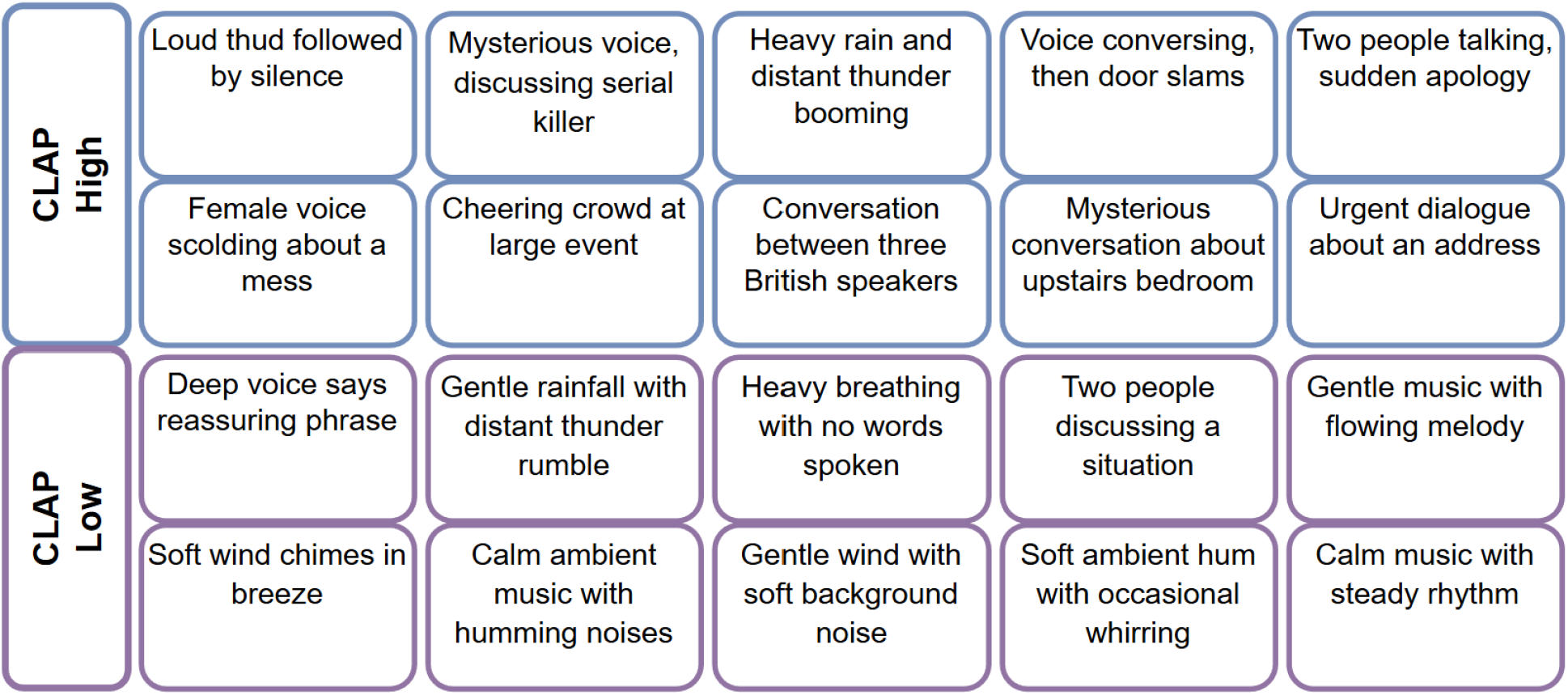
Short summarize by (GPT-audio; OpenAI, 2025) in audio clip that scored maximum and minimum value for PC2 CLAP predictions

**Figure S3.**
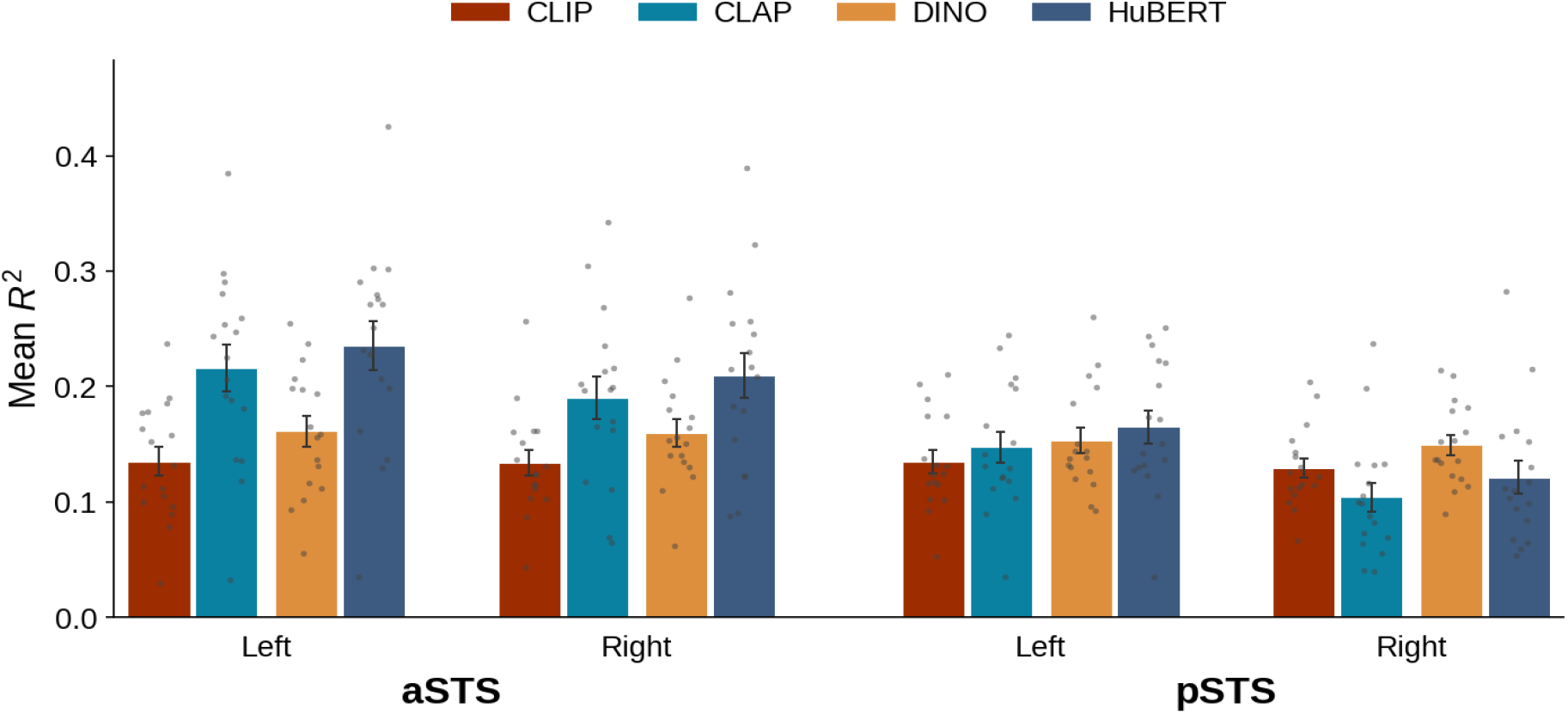
Neural response prediction by language-aligned and self-supervised encoders across STS ROIs. Mean R² (150 randomly selected voxels per hemisphere) for four embedding models: CLIP Vision and DINOv2 (visual), CLAP Audio and HuBERT (auditory), in anterior and posterior STS. Results show the mean across 17 participants; dots show individual participants. Within each modality, the self-supervised, non-language-aligned encoders (DINOv2, HuBERT) perform better than their language-aligned counterparts (CLIP, CLAP), with HuBERT numerically highest in anterior STS. This might suggest that STS prediction is driven by the richness of the learned high-level representations rather than by language supervision specifically.

**Figure S4.**
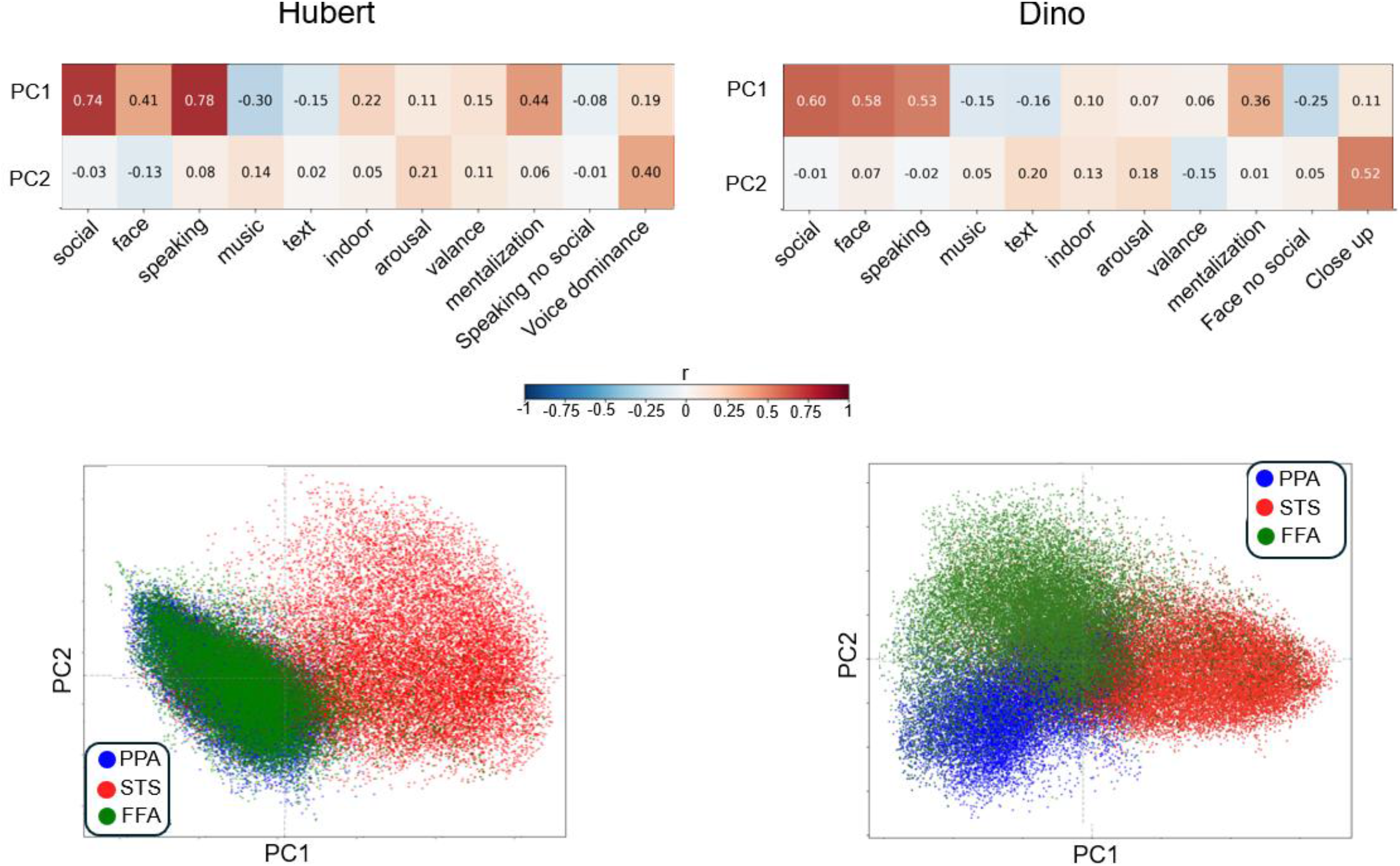
Principal components capture social-affective-perceptual information represented in neural responses for Dinov2 and Hubert, same pattern as for language supervised models (CLIP and CLAP). Loadings of each voxel on each PC. Colors represent different functional ROIs (PPA, STS, FFA).

**Fig S5.**
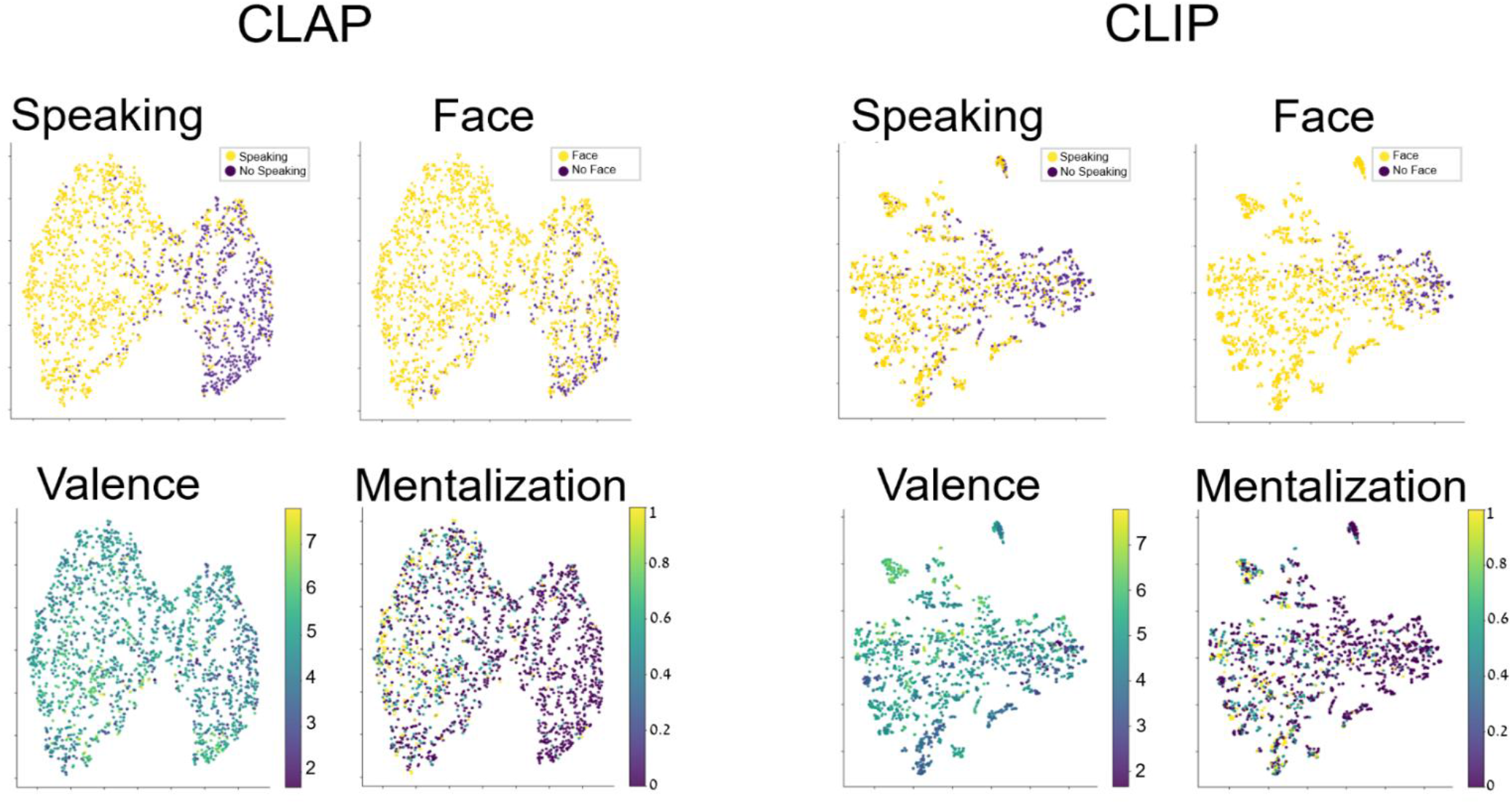
Geometric structure of predicted STS responses. t-SNE visualization of predicted activity for CLAP (left) and CLIP (right) encoding models for each movie frame and sound, color-coded by annotations based on human manual annotations. We included voxels from the EBA and ran the same PCA analysis described in the main text. Results reveal the same first PC1 that reflects social interaction in visual and audio input and was localized to the STS. CLIP PC2 was moderately correlated with face and was localized in the EBA. CLIP PC3 was similar to PC2 in the model that did not include EBA, corresponding to shot type (short-long). CLAP PC2 and PC3 capture voice dominance.

**Figure S6.**
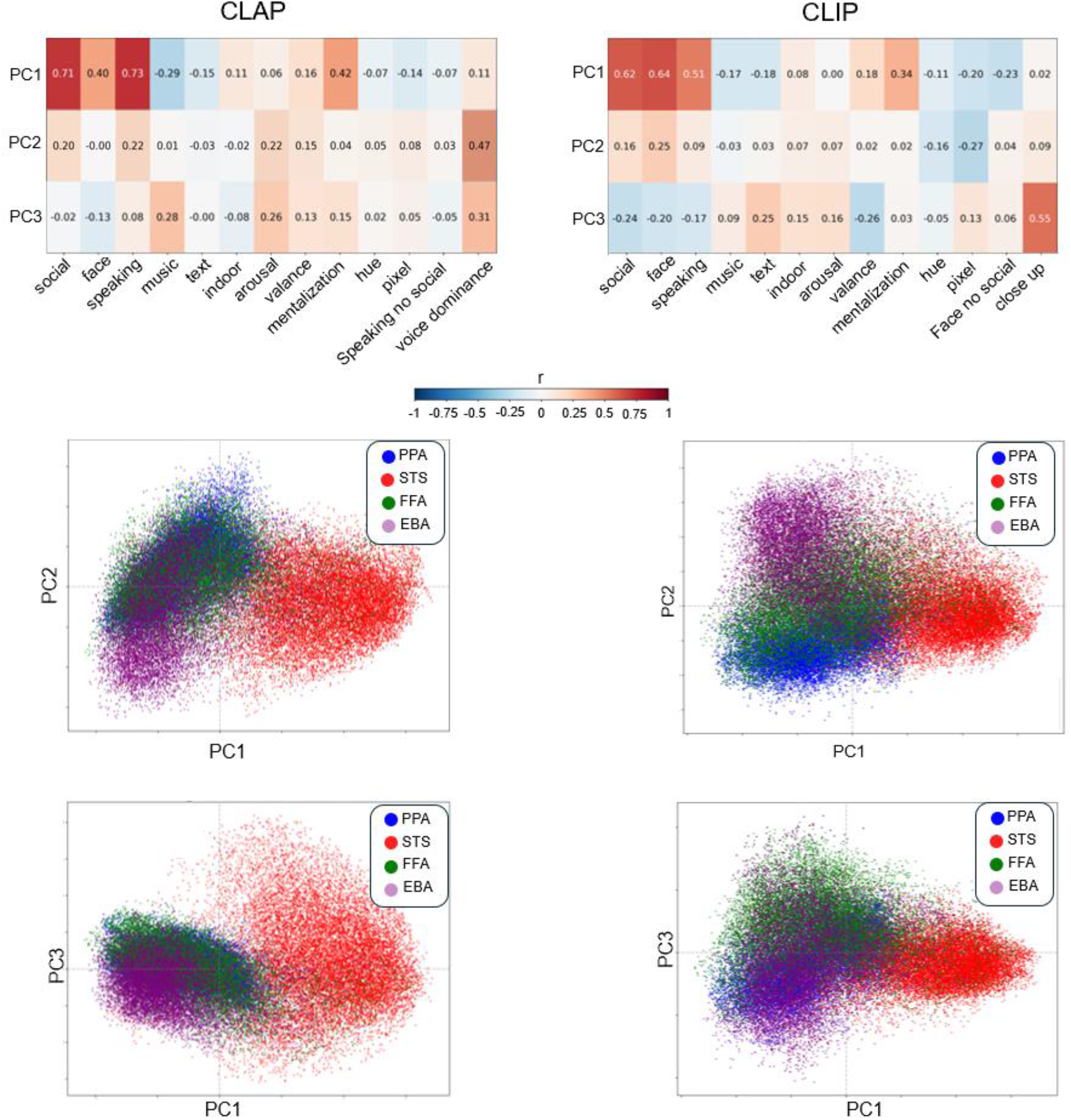
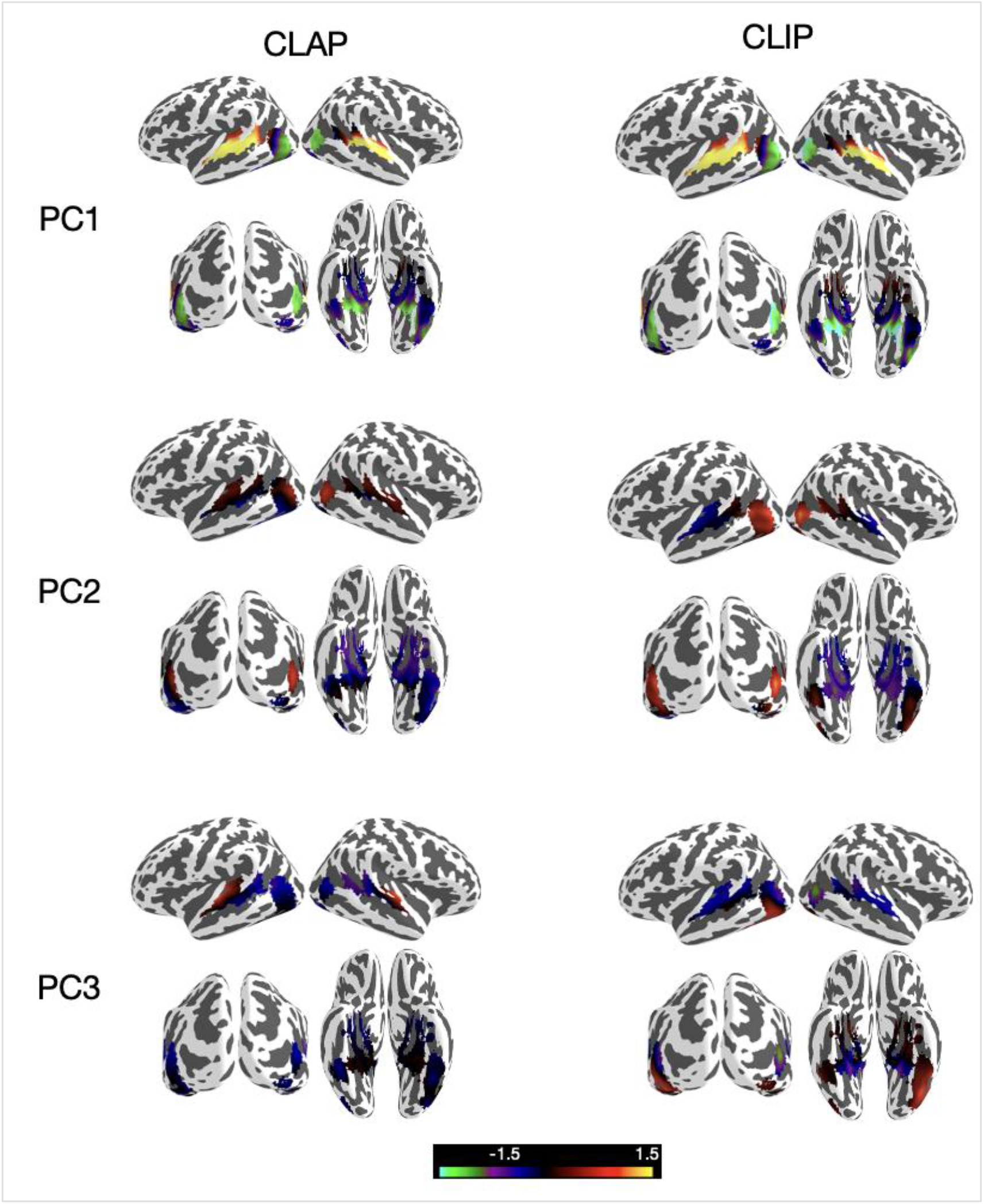
Top panel: the correlations between principal components on the encoding model of voxels in the PPA, STS, FFA, EBA and human annotated social-affective features for CLAP (left) and CLIP (right). Middle panel: Loadings of each voxel on each PC. Colors represent different functional ROIs. Bottom panel: PC loading scores projected onto inflated cortical surfaces for CLAP (left) and CLIP (right) for PC1, PC2, and PC3. Warm colors indicate positive loadings; cool colors indicate negative loadings (scale: −1.5 to 1.5).

